# MISATO - Machine learning dataset of protein-ligand complexes for structure-based drug discovery

**DOI:** 10.1101/2023.05.24.542082

**Authors:** Till Siebenmorgen, Filipe Menezes, Sabrina Benassou, Erinc Merdivan, Stefan Kesselheim, Marie Piraud, Fabian J. Theis, Michael Sattler, Grzegorz M. Popowicz

## Abstract

Large language models (LLMs) have greatly enhanced our ability to understand biology and chemistry. Yet, relatively few robust methods have been reported for structure-based drug discovery. Highly precise biomolecule-ligand interaction datasets are urgently needed in particular for LLMs, that require extensive training data. We present MISATO, the first dataset that combines quantum mechanics properties of small molecules and associated molecular dynamics simulations of about 20000 experimental protein-ligand complexes. Starting from the PDBbind dataset, semi-empirical quantum mechanics was used to systematically refine these structures. The largest collection to date of molecular dynamics traces of protein-ligand complexes in explicit water are included, accumulating to 170 μs. We give ML baseline models and simple Python data loaders, and aim to foster a thriving community around MISATO (https://github.com/t7morgen/misato-dataset). An easy entry point for ML experts is provided without the need of deep domain expertise to enable the next generation of drug discovery AI models.

---

In recent years, Artificial Intelligence (AI) predictions have revolutionized many fields of science. In structural biology, AlphaFold 2^1^ predicts accurate protein structures from amino acid sequences only. Its accuracy nears state-of-the-art experimental data. The success of AlphaFold 2 is made possible due to a rich database of nearly 200 000 protein structures that have been deposited and are available in the protein data bank (PDB)^2^. These structures were determined over the past decades using X-ray crystallography, Nuclear Magnetic Resonance (NMR), or Cryo-Electron Microscopy (Cryo-EM).

Despite enormous investments, there are still few new drugs approved yearly, with development costs reaching several billion dollars^3,4^. An ongoing grand challenge is rational, structure-based drug discovery (DD). In contrast to protein structure prediction, this task is substantially more difficult.

At early stages of DD, structure-based methods are popular and efficient approaches. The biomolecule provides the starting point for rational ligand search. Later, it guides its optimization in order to optimally explore the chemical combinatorial space^5^ while still ensuring drug-like properties. *In-silico* methods that are in principle able to tackle structure-based DD include semi-empirical Quantum Mechanical (QM) methods^6^, Molecular Dynamics (MD) simulations^7,8^ docking^9^, and coarse grained simulations^10^, which can also be combined to be more efficient. Yet, these methods suffer from either generally low precision or are computationally too expensive while still requiring substantial experimental validation. Recent examples show that classical, ball-and stick atomistic model representations of biomolecular structures might be too inaccurate in certain situations to allow for correct predictions^11–14^.

Introduction of AI into the process is still at an early stage. AI approaches are, in principle, able to learn the fundamental state variables that describe experimental data^15^. Thus, they are likely to abstract from electronic and force field-based descriptions of the protein-ligand complex. Yet, so far mostly simple solutions have been proposed that do not incorporate the available protein-ligand data to their full extent, like scoring protein-ligand Gibbs free energies^16,17^, ADME property estimation^18^, or prediction of synthetic routes^15,19,20^. Most of these approaches are constructed using one-dimensional SMILES^21,22^ and only few attempts have been made to properly tackle 3D biomolecule-ligand data^23–28^.

Due to the recent success of LLMs^29^, these models are starting to get introduced into the drug discovery process^30–33^.

Several databases are available that contain experimental structures of protein-ligand complexes, usually extracted from the PDB (*e*.*g*., PDBbind^34^, bindingDB^35^, bindingMOAD^36^, Sperrylite^37^). Only recently the first database of MD derived traces of 5000 protein-ligand structures was reported^38^. In spite of these efforts, so far no AI model has been proposed that convincingly addresses the rational drug discovery challenge in the way that AlphaFold 2 answered the protein structure prediction problem^39,40^.

The current structure-based AI models are severely hindered by several factors: neglecting the conformational flexibility (dynamics and induced fit upon binding); entropic considerations; inaccuracies in the deposited structural data (incorrect atom types due to missing hydrogen atoms, incorrect evaluation of functional group flexibility, inconsistent geometry restraints, fitting errors); chemical complexity (*e*.*g*., non-obvious protonation states); overly simplified atomic properties; highly complex energy landscapes in molecular recognition by their targets. Attempts to train AI models currently require to infer this missing information implicitly. Yet, with a limited number of publicly available protein-ligand structures (ca. 20000) and lack of thermodynamic data, this inference is failing. Thus, it is preventing structure-based models from producing groundbreaking results^39,40^.

Here, we propose a new protein-ligand structural database MISATO (**M**olecular **I**nteraction**S A**re struc**T**urally **O**ptimized) that is based on experimental protein-ligand structures. We provide a quantum chemical-based structural curation and refinement, including regularization of the ligand geometry. We augment this database with missing dynamic and chemical information, including molecular dynamics in a timescale allowing the detection of transient and cryptic states for certain systems. The latter are very important for successful drug design^41^. Thus, we supplement experimental data with the maximum number of physical parameters. This eases the burden on AI models to implicitly learn all this information, allowing focus on the main learning task. The MISATO database provides a user-friendly format that can be directly imported into Machine Learning (ML) codes without additional conversion. We also provide various pre-processing scripts to filter and visualize the dataset. Example AI baseline models are supplied for the calculation of quantum chemical properties (chemical hardness and electron affinity) and for the prediction of protein flexibility or induced-fit features (adaptability) to simplify adoption. We wish to transform MISATO into an ambitious community project with vast implications for the whole field of drug discovery.

## Results

### MISATO dataset

The basis for MISATO (Fig. 1) are the 19443 protein-ligand structures from PDBbind^34^. These structures were experimentally determined over the last decades and represent a diverse set of protein-ligand complexes for which experimental affinities are available. In the context of AI for drug discovery it is of utmost importance to train the models on a dataset with highest possible correctness and consistency, for a number of reasons: First, the total number of available structures is much lower compared to typical training sizes of other AI targets. Second, ligand association has a quite complex energy landscape during molecular recognition. Delicate deviations in the protein-ligand structures or atomic parameters can drastically impair binding. In the PDB, incorrect atom assignments and inconsistent geometries are not uncommon. More severe, hydrogen atoms are highly sensitive to their chemical and molecular environment and are rarely experimentally accessible. All these issues have been systematically addressed in our work and are compiled in our new database (Fig. 2 and 3).

**Fig. 1:**
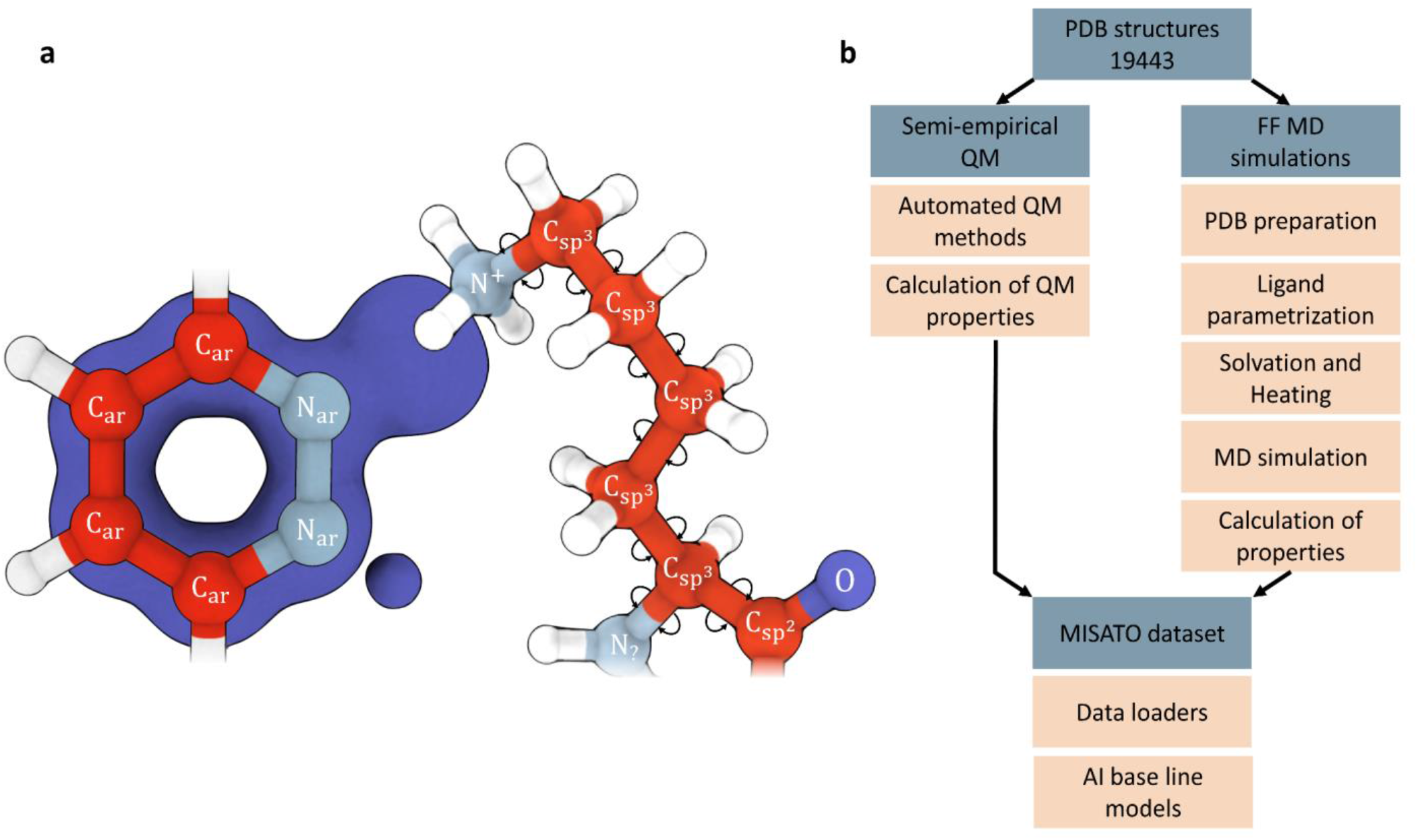
MISATO combines QM data with MD derived protein-ligand dynamics. **a**, We provide the first dataset that combines semi-empirical QM properties of small molecules with MD simulated dynamics of the entire, experimental protein-ligand complexes. All common errors in protein and ligand nomenclature, protonation, geometry, etc. are fixed. **b**, An overview of the dataset and the applied protocols including data preparation, preprocessing and AI base line models is given.

**Fig. 2:**
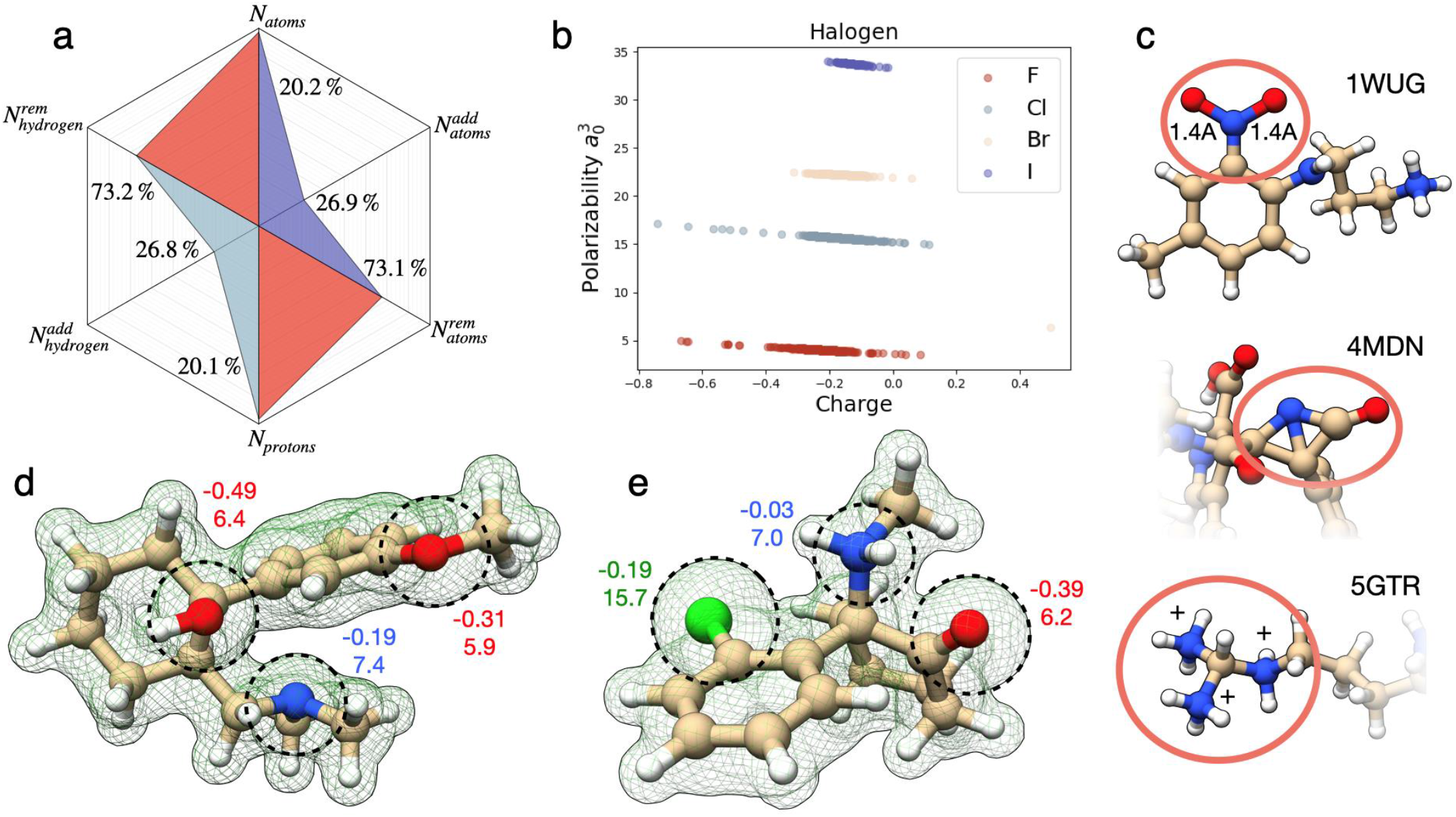
Changes applied to the PDBbind database based on our quantum chemical protocol. **a**, Statistical overview of changes introduced by our optimization protocol. *N*_*atoms*_ corresponds to total changes in the atom count compared to the source database. In most cases atoms were removed 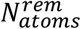); only in 27 % of the cases the number of atoms was increased, 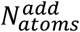. Similar considerations apply to protons - light blue; *N*_*protons*_. **b**, D4-Polarizability Vs. Partial Charge for all the halogens in database. The outliers were analyzed to find possible wrong atom assignments. This was the case of the bromine atom in the lower right corner, which in reality is a boron.**c**, Examples of inconsistent structures: 1WUG contains overly elongated NO bonds; 4MDN contains a nitrogen in angular violation of VSEPR; 5GTR shows a typical problem in the protonation state.**d, e**, Calculated electronic density for ketamine (4G8H) and tramadol, respectively. Dashed circles show the size of electronic density around selected atoms. The numbers next to those atoms represent partial charge (up) and atomic polarizability (down). These are electronic descriptors representing the electronic density around each center.

**Fig. 3:**
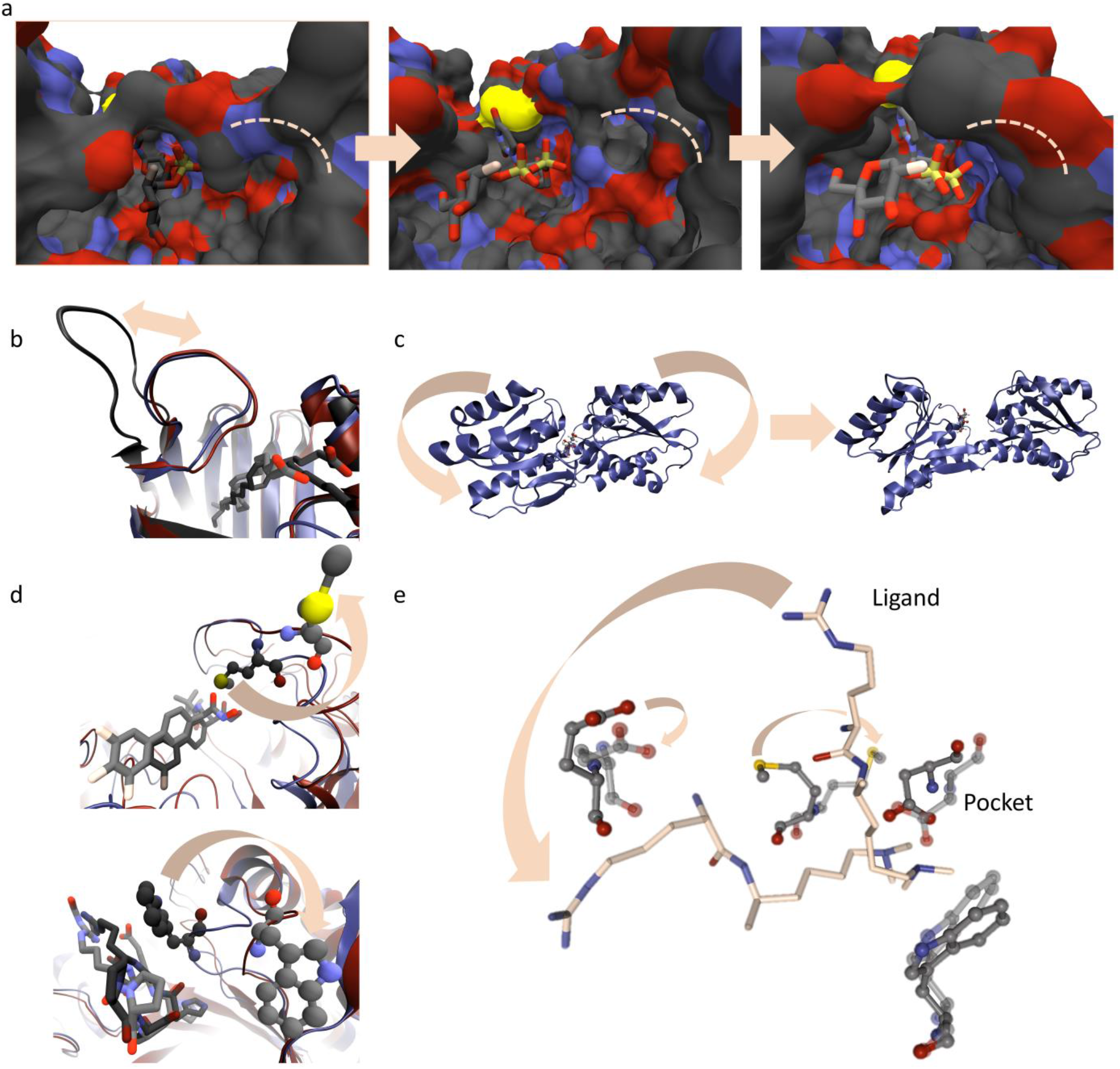
Overview of events captured by the MD simulations in the binding pocket. **a**,**b**,**c** Reversible opening and closing of the binding pocket can be captured during the simulations, including cryptic binding sites. The structure of 2AM4 is shown after 2ns (left panel), 6ns (middle panel) and 10ns (right panel) simulation time (**a**). The opening loop region (**b**,structure 2LKK) is visualized for superimposed timesteps (2ns: blue cartoon, dark hue, 6ns: black cartoon, medium dark hue, 10ns: red cartoon, light hue). The protein pocket opens in structure 8ABP during the simulation (**c**). **d**, Protein residues at the binding site can undergo large adaptations within the simulations, indicating unstable interactions or possible switches. This is shown for a Methionine residue for 4ZYZ (upper panel) and a Tryptophane residue of 1WAW (lower panel). Coloring as in b after 2ns and 10 ns. **f**, MD simulations captured local adaptability of the binding pocket and ligand. For example, in structure 2IG0 parts of the ligand (licorice, carbons in ivory) are quite flexible in the protein pocket (gray carbons), when comparing the first (dark hue) and the last frame (light hue) of the MD run.

MISATO is publicly accessible and can be downloaded from Zenodo. We provide instructions for usage, data loaders via our github repository, and a container image with all relevant packages installed for GPU usage (see Table 1). The dataset is accessible via a Python interface using a simple *pytorch* data loader. Special attention was given to code modularity, which makes it easy to adjust the AI architecture (Fig. 4). We have implemented our dataset according to atom3d^42^ code base, a comprehensive suite of Machine Learning methods in the context of molecular applications.

**Table 1:**
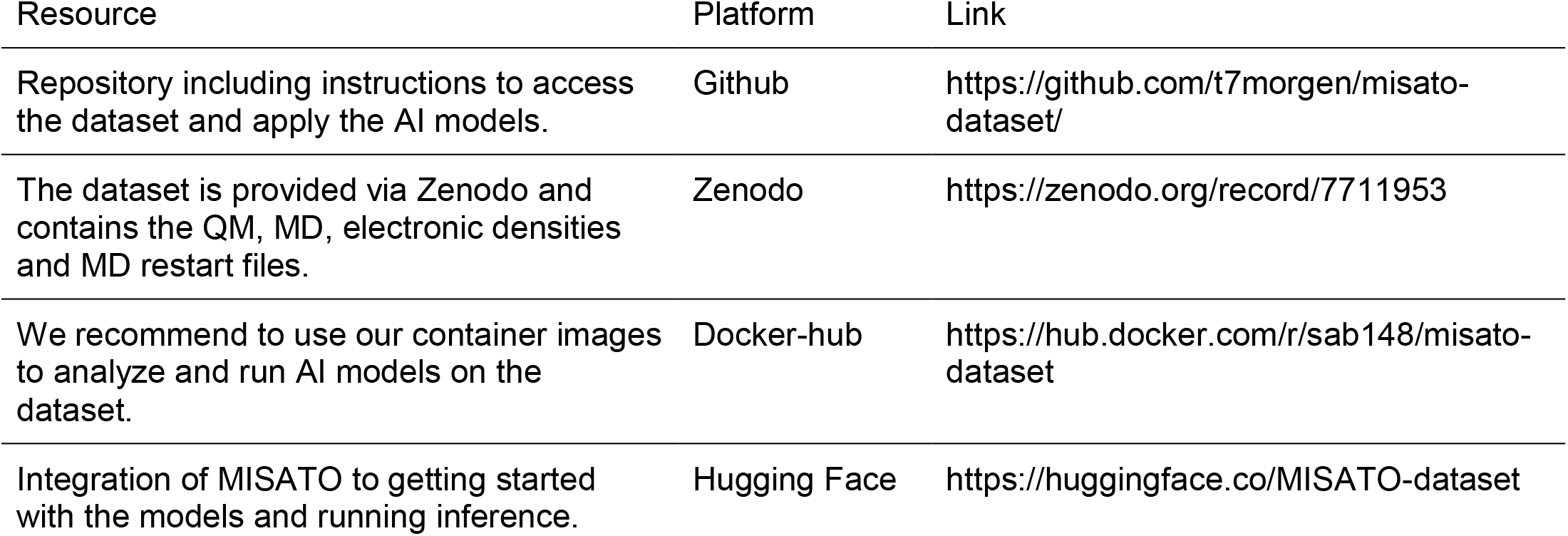
Overview of resources provided by the MISATO database.

| Resource | Platform | Link |
| --- | --- | --- |
| Repository including instructions to access the dataset and apply the AI models. | Github | https://github.com/t7morgen/misato-dataset/ |
| The dataset is provided via Zenodo and contains the QM, MD, electronic densities and MD restart files. | Zenodo | https://zenodo.org/record/7711953 |
| We recommend to use our container images to analyze and run AI models on the dataset. | Docker-hub | https://hub.docker.com/r/sab148/misato-dataset |
| Integration of MISATO to getting started with the models and running inference. | Hugging Face | https://huggingface.co/MISATO-dataset |

**Fig. 4:**
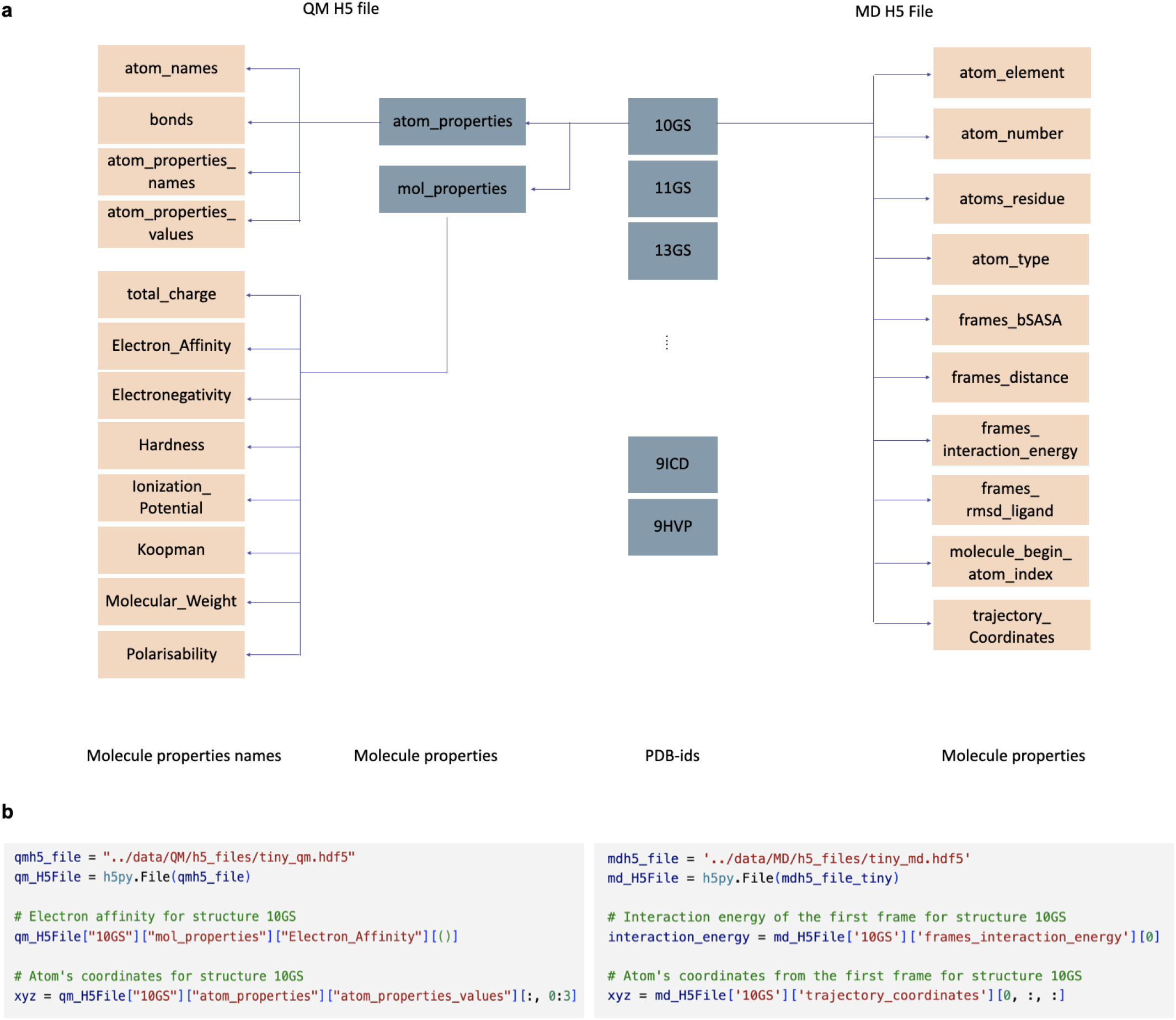
Data hierarchy of the QM and MD files. **a**, The QM data can be accessed via the PDB-id. The properties are split by atom properties and molecular properties. Examples of the calculated molecular properties are given. The electronic densities are provided in a separate file. **b**, The MD data is also subdivided by PDB-id. The properties are either calculated for all atoms, for each timestep (frame) or the whole trajectory, as indicated by the name. **c**, Example code to access the dataset files and the corresponding data loaders.

### Typical limitations in structural datasets

Understanding the nature and sources of errors in structural databases is imperative for improving the quality of the underlying molecular models. As MISATO is founded on experimental data, the two main sources of inaccuracies are limited spatial resolution of the experimental structures and problems associated with the software used for processing the molecular geometries. Besides absence of hydrogen atoms in crystallographic structures, resolution affects the heteroatom geometry. Contracted or elongated bonds are common (Fig. 2). For instance, most nitro groups we examined were heavily distorted: In 1WUG structure^43^, NO bonds are almost 17 % larger than reference experimental data^44^. Another example is seen in the 4MDN structure^45^, where an amide was so distorted that it explicitly violated VSEPR (valence shell electron pair repulsion) theory. Re-inspection of the experimental electronic density hinted that the CÔC angle in the 4-Chlorobenzyl phenyl ether moiety is also larger by almost 20º against anisole, a reference compound for that bond angle^44^. Simultaneous relaxation of both groups leads to significant improvement, in particular an amide group very close to reference structural values. Such errors in the heteroatom skeleton propagate further when assigning and counting hydrogen atoms. In the 5GTR structure^46^, a guanidino group strongly deviates from the expected planarity. The immediate consequences are incorrect atomic hybridizations and over-assignment of hydrogen atoms, with a local formal charge of +3 in a radius of one bond around the central carbon.

Many other issues are subtler. In 4IQT^47^ and 6FNE^48^, fine structural deformations coupled with infrequent functional groups led the proton assignment algorithms to miss an enol and an enolate and propose instead an alcohol and a ketone. From a chemical point of view, an enolate is a nucleophile, while ketones behave typically as electrophiles. This may be further followed in the respective dynamic traces since in those two simulations the ligands simply dissociate. In other cases, the protonation states are perfectly valid in an aqueous environment, however, highly unlikely in the respective protein’s pocket. 4MHZ structure^49^ would be a typical example, where, when binding takes place, an imidazole ring finds itself surrounded by 3 carboxylates. Such fine cases can only be identified when looking at the protein-ligand complex itself. Finally, we also observed chemically and biologically inconsistent data. PDBbind and PDB report the ligand in 2JJR^50^ to be an extremely powerful binder to a mutant of trichosanthin. Visual inspection of the complex reveals the ligand is a crystallization buffer: tris(hydroxyethyl)aminomethane (TRIS). Careful examination of the original reference gives away that the binding assay used to obtain the nanoMolar affinity of TRIS to trichosanthin is indeed unrelated to the proposed ligand. In other situations, the binding pocket proposed was incorrect. The 5OS8 structure^51^ shows one binding pocket and two additional adsorption points to the protein’s surface. Unfortunately, the coordinates selected to characterize the pair were for one of the adsorbing species.

### QM curation of ligand space

Consistent atomic assignments were determined using a series of semi-empirical tests. Semi-empirical quantum chemical methods offer a good compromise between accuracy and computational efficiency^52^, which is suitable to refine a collection of almost 20000 structures of various chemical nature and dimensions (from 6 to almost 370 atoms per molecule). The consistency tests we designed were performed in vacuum to ensure maximum sensitivity of the calculations to structural inconsistencies. Predicted properties, however, are also obtained using implicit solvation.

It is well documented that molecules with many polar groups lack convergence in wavefunction optimization^53^. The same applies when incorrect charges or protonation states are used. Implicit solvation significantly ameliorates the issue and masks problems. In fact, after determining the first guess for total molecular charges, single-point-energy calculations on unrefined ligands using implicit water required roughly 6 hours of computation time. Turning off implicit solvation increased the calculation time to almost three weeks on the same machine. This was indicative of severe limitations in proton and total charge assignment. Alternative protonation algorithms were tested, *e*.*g*., OpenBabel^54^. Due to experimental inaccuracies in the geometries, results were still faulty (SI Fig. S4-S7).

Our refinement protocol started with a search for structures with strong atomic overlap. Next, we looked for structures with problematic wavefunction convergence. Vanishing HOMO-LUMO gaps or unpaired electrons flagged further problems, just like violations of the octet rule based on QM population analysis. Lastly, we searched for changes in ligand connectivity patterns after QM geometry optimization. This was particularly useful in determining inconsistent protonation states or incorrect electron counting, which generated biradicals. Calculated properties yielded additional testing grounds. Incorrect element assignments were detected when plotting the partial charges against D4 polarizabilities^55^ (Fig. 2 b).

Severe structural deformations were also detected, inconsistent with the chemical structure (see previous section). For the current stage of the database we decided to fix only the most extreme cases. This was done using Avogadro (SI Fig. S2)^56^. Further structural refinement is planned using experimentally constrained ligand optimization.

Whenever our corrections would seem questionable, or the structure was unclear, we checked the original publication. Oxidation states were another sensible point for ligands containing transition metals. Examples of structures we refined are given in the SI (Fig. S1, S2, S3). To ease the inclusion and processing of new structures, a heuristics-based program is included in the database, which performs the basic structural processing (SI for more detail).

### Evaluation of the QM-based ligand curation

Employing the protocol defined in the previous section we modified a total of 3930 structures, which corresponds roughly to 20 % of the original database (Fig. 2). 3905 cases involve changes in protonation states, while changes in heteroatoms involves 97 ligands. These are predominantly the addition of model functional groups to emulate covalent binding with the protein (20) or the addition of missing hydroxyl groups to boronic acids.

Some ligands were split in several molecules as the original structures were not binary protein-ligand complexes (1 ligand): 1A0T, 1G42, 1G9D, 2L65, 3D4F, and 4MNV. 1E55 is supposed to be a mixture of two entities. However, the closest contact between them is insufficient to consider them separately, but also too large for a covalent interaction. Similar considerations apply to 1F4Y, though here it seems that close intramolecular contacts are at stake. In 4AW8 we observed a significant deformation for the published ligand, PG6. On close literature inspection, we observed that the reference affinity is related to the metal ion in the system, Zn(II), and not to PG6. The structure was consequently excluded.

As may be followed in Fig. 2, the most common adjustment was the removal of (hydrogen) atoms from the initial PDBbind geometry. This amounts to almost 75 % of the modifications. It has been pointed out that libraries like PDBbind possess biased datasets in terms of binding configurations^39^. The problems we have discussed thus far have been, to the best of our knowledge, so far never addressed.

### QM derived properties

We calculated several molecular and atomic properties for the ligands (SI Tab. S1). For the former, we include electron affinities, chemical hardness, electronegativity, ionization potentials (by definition and using Koopman’s theorem), static logP, and polarizabilities. The latter were obtained in vacuum, water, and in wet octanol. Atomic properties include partial charges from different models, atomic polarizabilities, bond orders, atomic hybridizations, orbital- and charge-based reactivity (Fukui) indices, and atomic softness. Reactivity indices and atomic softness are derived for interactions with electrophiles, nucleophiles, and radicals. Finally, we also provide tight-binding electronic densities for all ligands. Partial charges were calculated at several levels, as these are somewhat method-sensitive quantities. AM1 charges are usually the starting point for charge correcting schemes to be used in molecular dynamics simulations. This is the case of AM1-BCC^57^. Taking our AM1-CM1 charges and multiplying them by 1.14 (in the case of neutral molecules) yields 1.14*CM1A-LBCC charges^58^ used in OPLS-AA simulations^59^. The main advantage of the charges we provide is that these were obtained, when required, with a HOMO-LUMO level-shift to ensure convergence to sensible electronic states. Beyond MD simulations, CMx charges^60–62^ have also been shown to provide good estimates of molecular dipole moments just like tight-binding Mulliken charges^63^. From the latter, we infer furthermore on the reasonability of the electronic densities provided.

Regarding ionization potentials and electron affinities, the situation becomes delicate, since most quantum chemical methods fail to predict reasonable values^64^. This applies not only to DFT but also to *ab initio*. In the SI we provide a parameter study (SI Tab. S3) performed with data collected from the CCCBDB database^44^, verifying the generality of trends reported in the literature^64,65^. The parameter study shows furthermore that semi-empirical ionization potentials are of similar quality, if not higher, to the best DFT results. The advantage, however, is that we systematically apply the same level of theory for all molecules, small and very large alike.

### MD simulations

Experimental structural data are static snapshots that are assumed to represent a thermodynamic most stable state trapped in a crystal, but ignore the presence of conformational dynamics. Experimental description of dynamics in biological macro-molecules from ns to ms timescales is challenging and requires a combination of different spectroscopic techniques. NMR spectroscopy and fluorescence-based methods can provide relevant information, but are time consuming and so far, the dynamic information is not well captured in public databases. Molecular dynamics simulations can be performed, starting from experimental structures and letting them evolve in time using a force field that describes the molecular potential energy surface. Typically, time spans of nano to micro-seconds can be achieved for individual systems, depending on system size. MD traces allow the analysis of small range structural fluctuations of the protein-ligand complex, but in some cases large-scale rare events can be observed (Fig. 3). In existing drug discovery software these events are mostly neglected. MD simulations of 16972 protein-ligand complexes in explicit water were performed for 10 ns. Structures were disregarded whenever non-standard ligand atoms or inconsistencies in the protein starting structures were encountered. A variety of metadata were generated from the simulations to facilitate future AI learning (Fig. 4, SI Tab. S1, Fig. S7). The *RMSD*_*ligand*_ (RMSD of the ligand after alignment of the protein) and the RMSD of the whole complex were calculated with respect to the native structure. Also, binding affinities were estimated using MMGBSA scoring (no entropic contributions explicitly considered)^66^. Moreover, the buried solvent accessible surface area (SASA) was obtained for the complex. Calculated properties are stable over the simulations, proving them well-equilibrated (Fig. S7). For some systems, larger rearrangements of the binding site were captured that in the extreme cases led to an opening of the whole binding pocket (Fig. 3). These rare events give an indication of possible cryptic pockets or transient binding modes. In a small fraction of cases dissociation was detected.

### AI models

To exemplify possible applications of our dataset, baseline AI models were trained and evaluated. These are included in the repository as a template for future community development. For the QM dataset, the electron affinity and the chemical hardness of the ligand molecules were predicted (Fig. 5). The Pearson correlation is about 0.75 for electron affinity and 0.77 for chemical hardness. The MAE shows close predictions to the target values: on average 0.12 eV for electron affinity and 0.13 eV for chemical hardness. For these two exemplary QM features, high accuracy was achieved, opening a route to a fast derivation of QM properties. This is particularly important for larger molecules, where long calculation times are frequent.

**Fig. 5:**
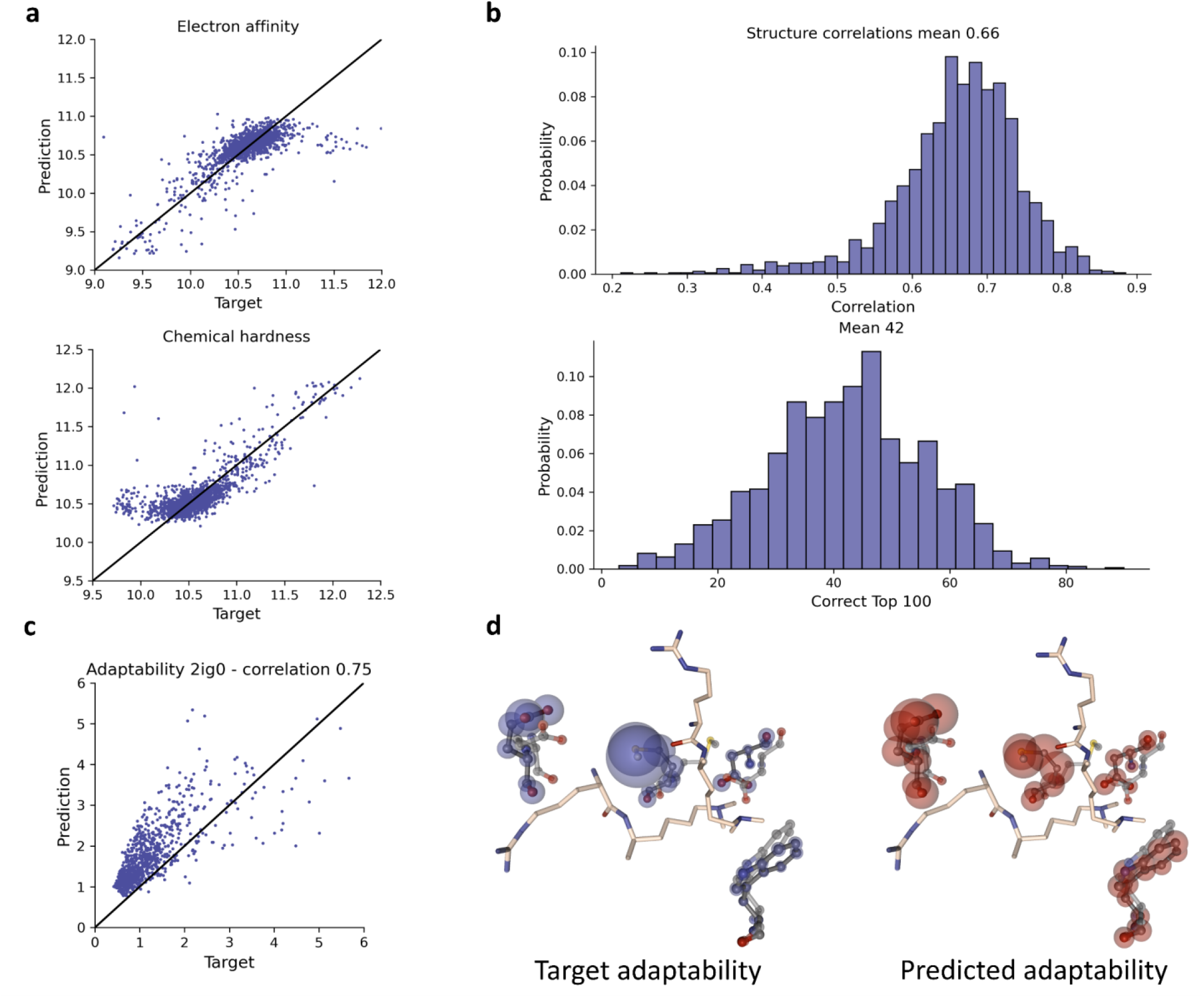
Performance of the AI baseline models. **a**, Scatter plot of the predicted against target values of chemical hardness and electron affinity. The AI baseline models to predict QM properties have a high correlation of 0.75 and 0.77 for electron affinity and chemical hardness, respectively. **b**, Adaptability is a measure of the per atom conformational plasticity of the protein. Histogram of the correlation and correct top 100 predictions of the adaptability for all structures in the test set are given. An overall mean correlation of 0.66 can be achieved and the mean top 100 accuracy was 0.42 for the adaptability predictions (MD). **c**, Scatterplot for the adaptability result (like in panel a) of example structure 2IG0. The predicted values are more narrowly distributed than the actual values but the general trend is correct, as shown by a high correlation value of 0.75. **d**, The adaptability of the residues in the protein pocket highly deviates between the amino acids. The AI model predicts the adaptability given in blue (target) and red (AI-predicted) spheres. The radius is scaled according to the adaptability value. The model is able to correctly identify the rigid residues (small spheres) but also the amino acids with high flexibility.

For the MD traces, the induced fit capability of the protein (adaptability) was predicted. The model was able to identify elements of biomolecule structure likely to adapt to ligand binding. We achieved a mean Pearson correlation of 0.66. On average 42 of the top 100 atoms were correctly predicted (Fig. 5). As given in Figure 5d, the model is capable to predict the atoms in the protein pocket that are mostly flexible during the MD run (large spheres) and also detect the more rigid protein regions (small spheres). This allows a fast examination of the protein pocket without the necessity of a lengthy MD setup and simulation. The adaptability model gives an innovative example of how experimental structures can be enhanced from the MD-based MISATO data.

## Discussion

The great advances over the last years of AI technologies were only possible due to huge datasets that are fed into these models. In structural biology, the protein folding problem was solved recently, but the drug discovery community still lacks a breakthrough model.

Here, we present MISATO, a database that will open novel routes in drug discovery. MISATO contains the first quantum-chemically refined ligand dataset, which permitted elimination of several structural inaccuracies and crystallographic artifacts. Our refinement-protocol can be immediately applied by others, for quick database augmentation. We enhance the curated dataset following two orthogonal dimensions. On the one hand, a quantum mechanical approach supplying systematic electronic properties. On the other hand, a classical approach that reveals the system’s dynamics and includes the binding affinity and conformational landscape. MISATO contains the largest collection of protein-ligand MD traces to date. Checkpoint files are made available for potential community extension of the dynamic traces (Table 1). Structural biology datasets until now are unable to incorporate entropy related information about binding sites and the dynamics of the systems. By conducting MD simulations, it is possible to approximate the conformational space for entropy estimation. A python interface, built to be intuitively used by anyone, provides pre-processing scripts and template notebooks.

The dataset augmentation presented here paves the route for creative applications of AI models. Possible metrics for the ML community are: docking, solvation, entropy, solubility, crystallization properties/conditions, protonation, matched molecular pairs generation, ligand growing, scaffold hopping and de novo generation. Our example GNN model offers quick access to pocket flexibility, a problem never tackled before. The adaptability prediction is a straight forward task for LLMs. This is however just a starting point for a whole new class of AI models sprouting from MISATO. Ultimately, we envision models building on the best of quantum and Newtonian worlds to obtain high quality thermodynamics, innovatively and efficiently matching the quality of experimental data. With MISATO, AI models will uncover hidden state variables describing protein-ligand complexes.

Together MISATO is meant to provide sufficient training power for accurate, next generation structure-based drug discovery using AI methods.

## Methods

### Semi-empirical calculations

QM calculations were performed using the ULYSSES library^67^, our in-house semi-empirical package. The methods of choice were GFN2-xTB^68^, AM1^69^ and PM6^70^. Implicit solvation was included using ALPB^71^ as parameterized for GFN2-xTB. Selected media included water and wet octanol. Bond orders and hybridizations were estimated using distance-based criteria. For further details on how the properties were calculated, please refer to the ULYSSES publication^67^.

### MD Simulations

For all MD simulations we used the Amber20^72^ software suite. The protein-ligand complexes were prepared and simulated based on a standard setup. We parameterized the ligands calculating AM1-BCC^57^ charges using antechamber^73^ (in case the charges did not converge within 1 hour we used AM1 charges calculated with ULYSSES). We used the gaff2^73^ force field for ligands and ff14SB^74^ for the proteins. The complexes were neutralized with Na^+^ and Cl^-^ ions and solvated in TIP3P^75^ explicit water using periodic boundary conditions in an octahedron box (minimum distance between protein and boundary 12 Å).

The complexes were minimized (1000 steps steepest descent followed by conjugate gradient) and heated to 300 K in several steps within 16 ps. We performed production simulations for 10 ns on all protein-ligand cases in an NVT ensemble. The first 2 ns were discarded as equilibration phase so that 8 ns are stored over 100 snapshots for each protein-ligand complex. Using pytraj^31^ we calculated different properties of the simulations like the MMGBSA interaction energy, the buried solvent accessible surface area (SASA), the center-of-mass (COM) distance between ligand and receptor and root-mean-square deviations (RMSD) from the native complex.

### Access to the database

The database can be downloaded from Zenodo (Table 1). Data is stored in a hierarchical data format (HDF). We created two H5 files, one for the protein-ligand dynamics and one for quantum chemical data, that can be accessed through our container images or after installation of the required python packages or via Hugging Face. Installation instructions are given in the repository (Table 1). Data is split for each structure using the PDB-id. The feature of interest must also be specified (Fig.4, SI Tab. S1). Python scripts are given in the repository showing how to pre-process the MD dataset for specific cases, only *C*_*α*_-atoms, no hydrogen atoms, only atoms from the binding pocket and inclusion of new features. Instructions how to run inference on new PDB files and visualize the baseline models are given. Checkpoint files for continuing the MD simulations and the electronic densities are provided separately.

### AI applications

For the baseline model for QM predictions we followed the GNN (Graph Neural Network) architecture for small molecule property prediction in atom3d^42^. This model is based on graph convolutions proposed by Kipf and Welling^76^ and was adapted for the simultaneous prediction of electron affinity and chemical hardness as essential parameters to describe the ligand. The performance of the ML model was evaluated using correlation and the mean absolute error (MAE).

We encode each molecule using the atom positions, the atom type, and the bond between the atoms. Each atom corresponds to a node. The atom types are one hot encoded and edges are defined by selecting the nearest neighbors with a distance of 4.5 Å for each atom. Edges are weighted inversely by the distance between the atoms. We removed outliers straying more than 20 standard deviations from the mean values (PDB-ids given in the SI). All outliers corresponded to molecules containing negatively charged groups and alkyl chains. In other words, these are highly saturated molecules from the electronic viewpoint. As a consequence of their electronic structure, acceptance of an electron is highly unlikely, resulting in very-low-to-negative electron affinities (EAs). Inaccuracies in the geometries further exacerbate the calculated EAs. The results on those systems indicate that some electronic properties are not quantitative, instead they simply reflect the system’s behavior. We trained the GNN with four NVIDIA A100 GPUs, 96 CPUs (from 48 physical cores) and for 200 epochs. We used a batch size of 128 and applied a random translation on each node of 0.05 Å.

For the MD task we modified the GNN architecture from atom3d^42^ for the node regression task by removing aggregation of node features into graph features. The architecture for the baseline model were five sequential GCNConv layers^76^ followed by two linear layers, summing to 370000 trainable parameters. The dataset was split into a train (80%), a test (10%) and a validation set (10%) (SI Tab. S2) by clustering the amino acid sequences of the proteins using Blastp^77^ in order to make sure to not have a leakage of similar structural motifs between the splits. We train the GNN with four NVIDIA A100 GPUs, 96 CPUs and for 15 epochs. We use a batch size of 8 and a random translation of 0.05 Å. With our model we calculated the adaptability of each atom during the MD simulation. To this end we performed an alignment of the coordinates of each simulation with reference to the first frame. In order to calculate the adaptability *γ*_*x*_ for each atom *x* we take the mean distance of each atom over all timesteps i to the initial position of the atom 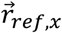:

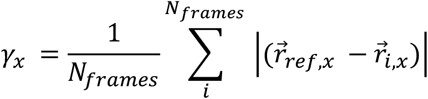

Hydrogen atoms were omitted to reduce the size of the model. For the evaluation the mean over the results for each individual structure was calculated. We evaluated the performance of our training using Pearson correlation and the average accuracy of the 100 most flexible atoms of each complex.

We used PyTorch version 1.14 to train the models. To code the data loaders and the GNN, we used PyTorch Geometric 2.3.0.

