## Supplemental_Information for "MISATO - Machine learning dataset of protein-ligand complexes for structure-based drug discovery"

### Supplementary Information

#### 1. Figures and Tables

**Tab. S1:** Details of the calculated QM properties for the ligand (left panel) and the MD properties on the dynamic traces (right panel).

| Ligand | Protein-ligand dynamics |
|---|---|
| Mulliken charges (AM1, PM6, GFN2-xTB, GFN2-xTB/water, GFN2-xTB/wet octanol) | MMGBSA interaction energy |
| AM1-CMx charges (x=1,2,3). | buried SASA |
| Atomic and molecular D4-polarizabilities (in gas, water and wet octanol) | COM-distance |
| Curated bond orders and atomic hybrizidations | rmsd complex |
| Molecular electronegativities | rmsd ligand |
| Molecular electron affinities | |
| Molecular ionization potentials (including Koopman) | |
| Molecular hardness | |
| Orbital and charge-based reactivity (Fukui) indices for electrophilicity, nucleophilicity and radical behavior | |
| Orbital and charge-based atomic softnesses with respect to electrophiles, nucleophiles and radicals | |
| static logP | |
| Electronic densities | |

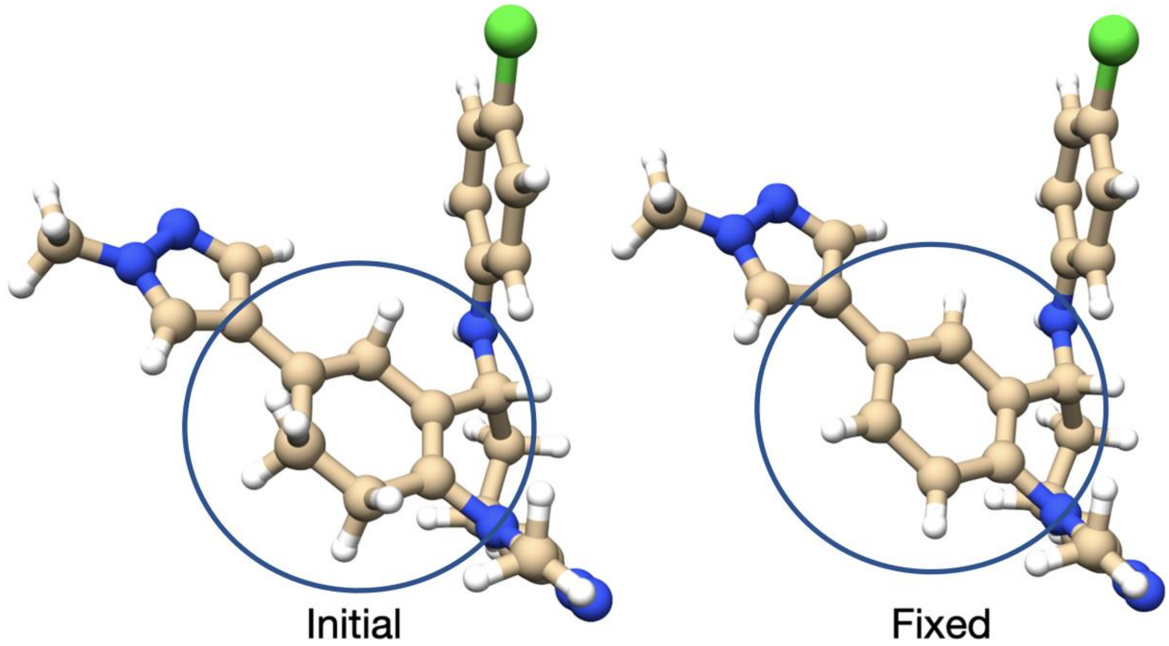


**Fig. S1:** Original and fixed structure for the ligand in 6K05. Inconsistencies in the geometries were identified using population analysis, which allowed us to further identify protonation states inconsistent with a given atomic hybridization.

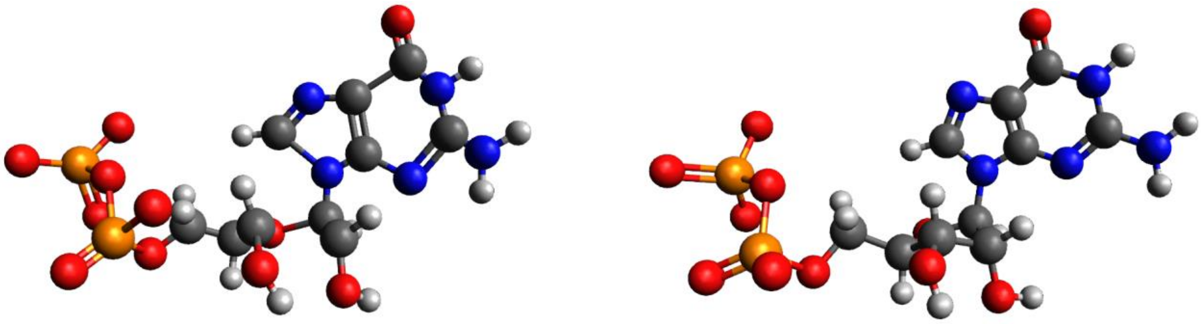

**Fig S2:** Original and fixed structure of the ligand 3ZY2 using Avogadro's UFF structure optimization tool.

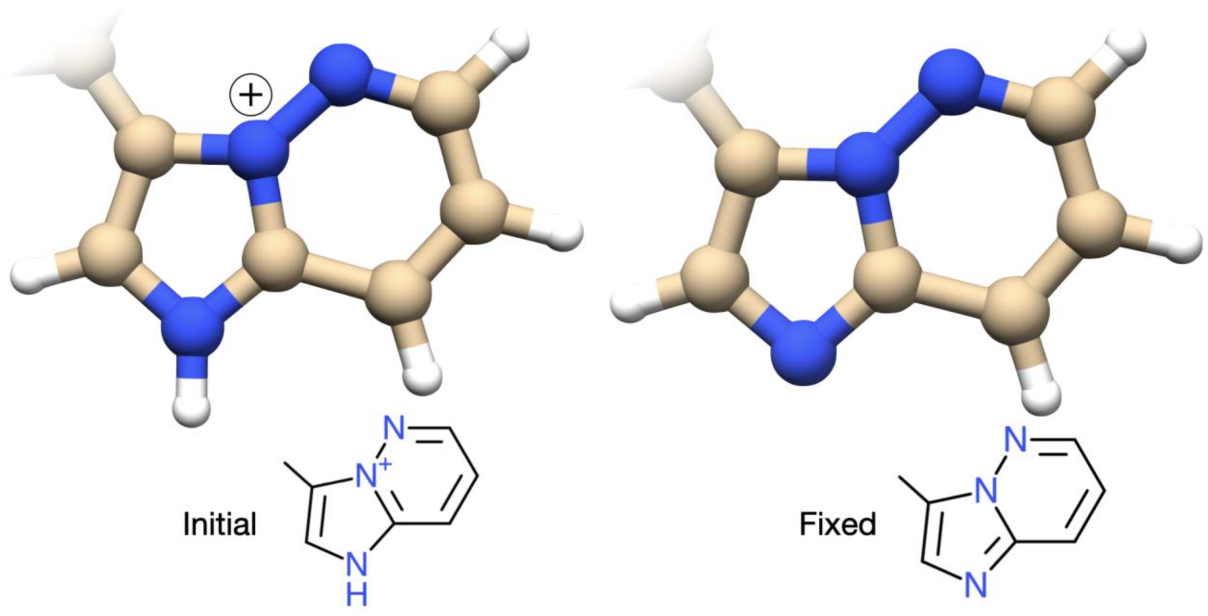


**Fig. S3:** Original and fixed protonation state for the ligand in the complex 1URW which contains an imidazo-[1,2-b]-pyridazine fragment. We applied the principle of charge neutrality. See 1PYE for another example.

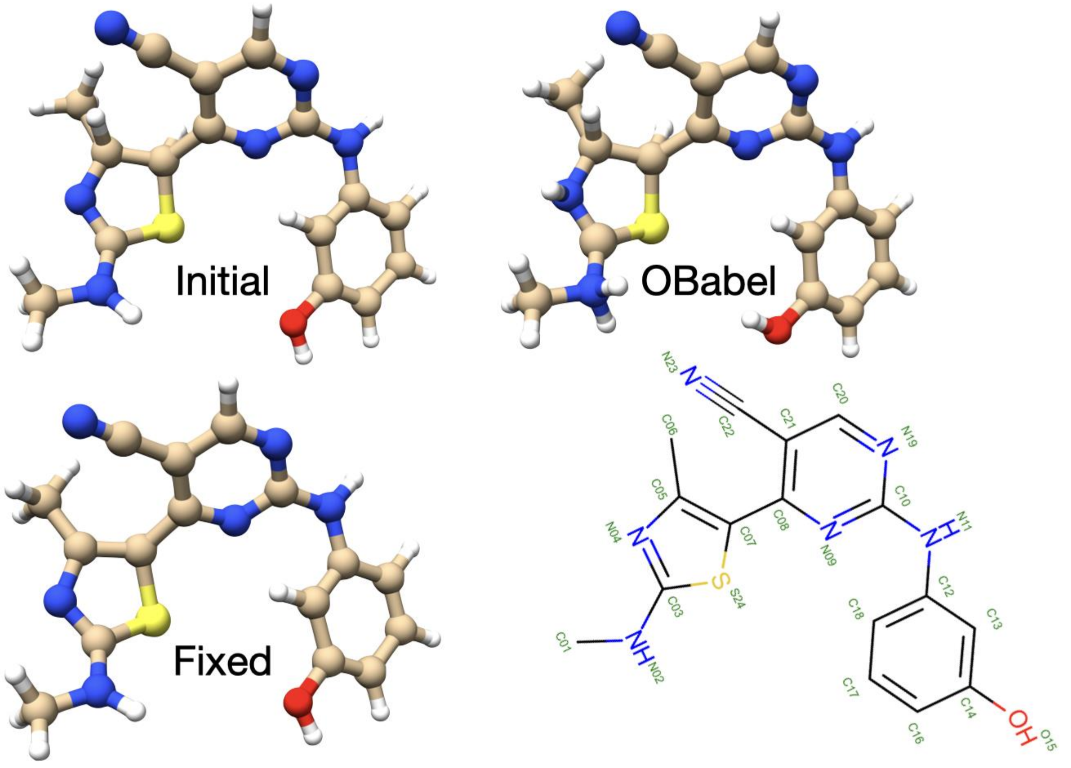


**Figure S4:** Initial, Openbabel and fixed structure for 4BCN. The two-dimensional representation of the structure reported in the PDB database is also given [put consistent 2D representations].

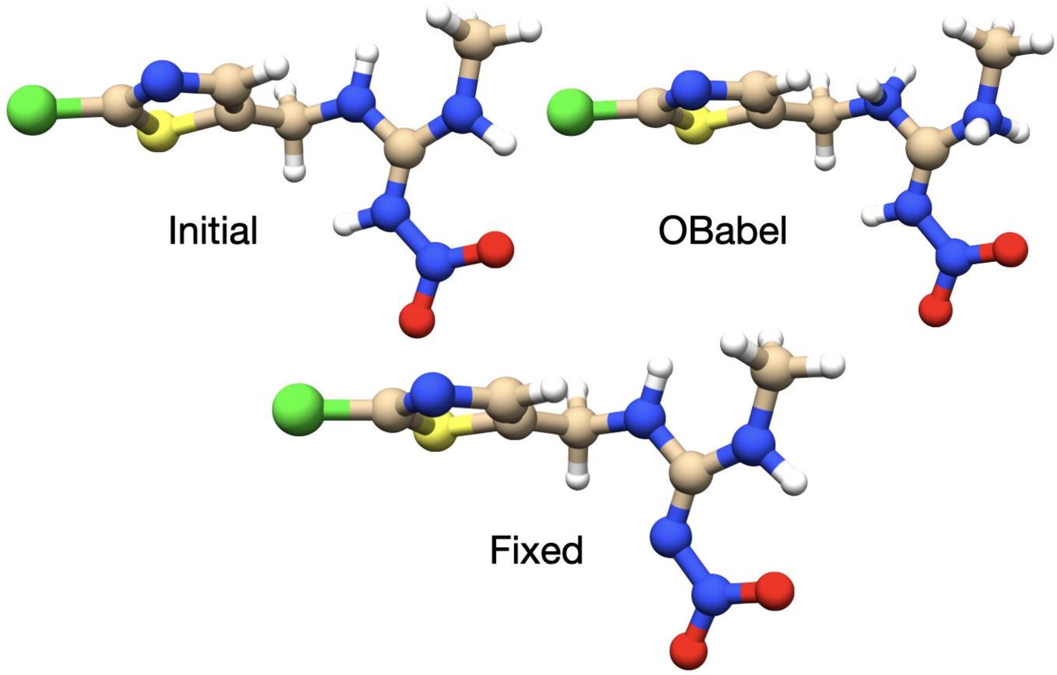


**Figure S5:** Initial, Openbabel and fixed structure for 2ZJV. Note that that it is highly unlikely that the nitrated nitrogen atom is protonated, due to mesomeric effects from the nitro group.

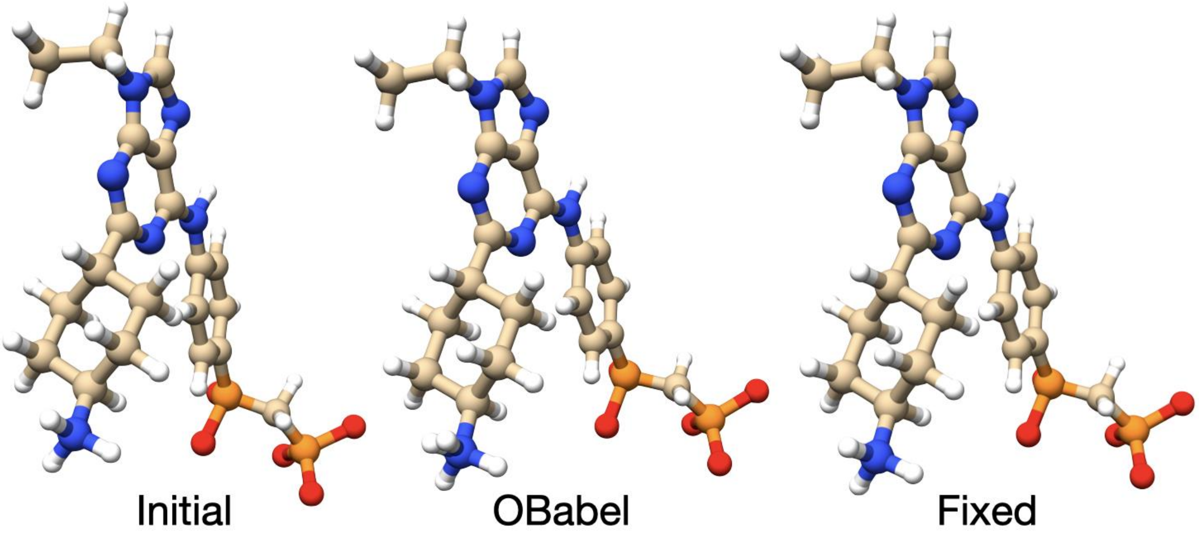


**Figure S6:** Initial, Openbabel and fixed structure for 2BDF. The lowest nitrogen atom (from amino group) is in explicit violation of the octet rule. Furthermore, there are two overlapping protons.

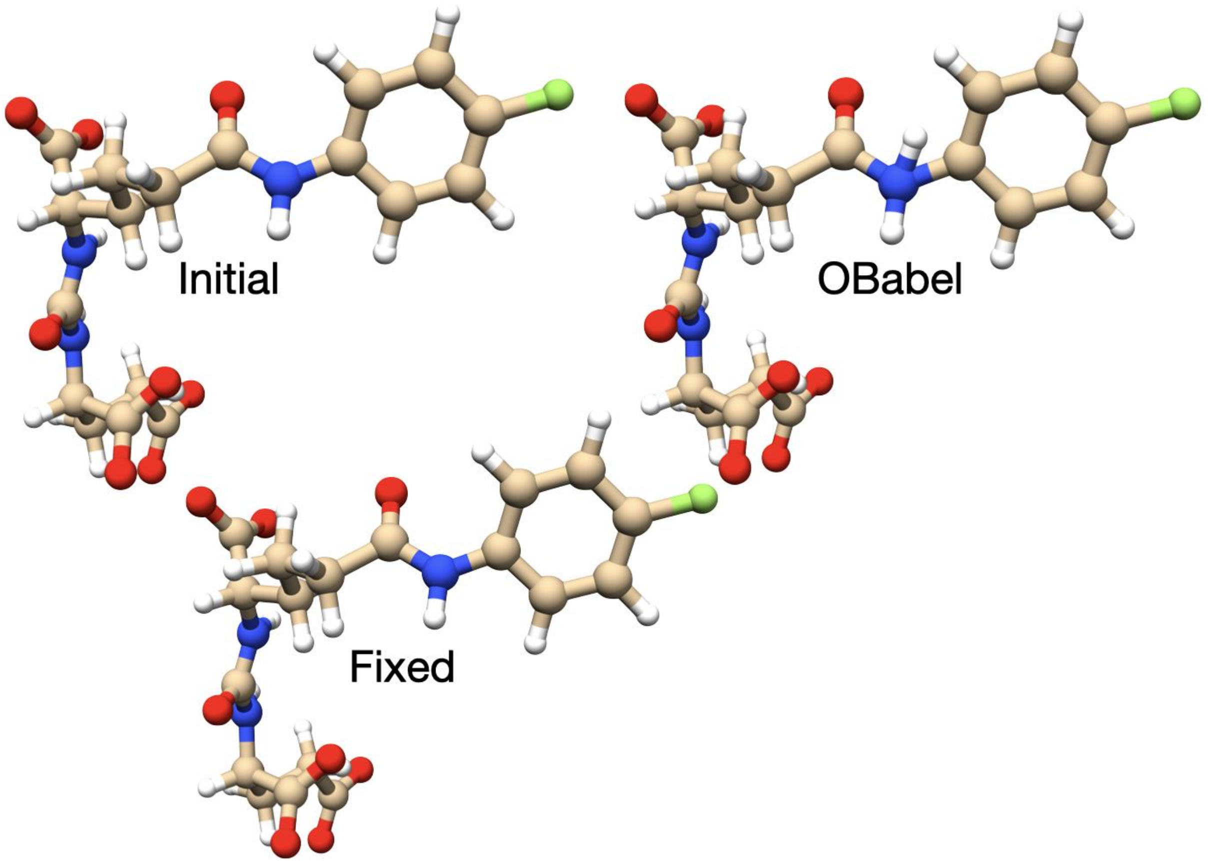


**Figure S7:** Initial, Openbabel and fixed structure for 6H7Y.

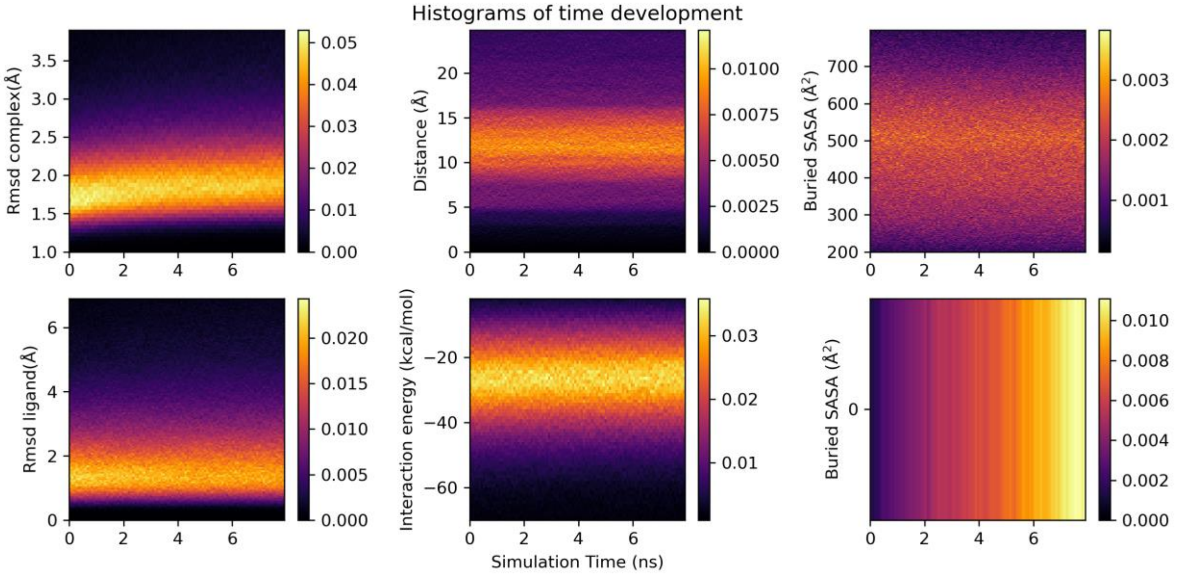


**Figure S7:** Heatmaps of the histograms of different properties of all protein-ligand complexes for each recorded timestep. The different colors represent different probabilities of a given bin from yellow (high probability) to magenta (low probability). In general, the calculated properties are quite stable through the simulation, which is a good indication that a converged ensemble is captured. Nevertheless, for individual cases the RMSD increases within the simulation time due to a conformational rearrangement of the structure.

**Tab. S2:** Number of samples in train, validation and test for each dataset presented in the paper.

| Task | Number of Samples | | |
|---|---|---|---|
| | Train | Val | Test |
| QM | 15506 | 1939 | 1938 |
| MD | 13765 | 1595 | 1612 |

#### 2. Added chemical groups

The list of structures in which we added a chemical group to model effects of covalent binding to the protein are

2FOU, 4JJG, 3CSL, 5WAD, 4XKC, 5OD5, 2P8O, 2Z97, 3W8O, 3RO0, 4Z46, 2FOY, 3ZS1, 2FOV, 5TYJ, 5TYK, 5TYL, 5TYN, 5TYO, 5TYP.

#### 3. Outliers not considered for the QM model

1YHM, 4U6C, 4DZW, 2HAW, 2Z50, 4DXJ, 5IJJ, 2ONB, 4E1E, 2IT4, 2RK8, 2O1C, 3T01, 3C14, 2F89, 2F94, 1A0TB, 1A5G, 4DGO, 4WM9, 6B1X, 1A46, 1A61, 2FSA, 4DWG, 1A0TA, 3BU8, 4UMJ, 3KXZ, 4HZX

#### 4. Evaluation of semi-empirical ionization potentials

Based on the data in the CCCBDB for Koopman ionization potentials, we constructed Table SIP. We note that though deviations may differ according to functional group, the panorama is generally the same: semi-empirical ionization potentials are of quality comparable to DFT ones, and in some occasions also superior or at least not inferior to MP2 charges. Note that ionization potentials show some dependence on basis set. For fairness in the comparison, we decided to stick to a single basis set of general use by the community of applied theoreticians. We also chose a fair basis set for the evaluation of the property.

#### 5. Heuristics based program for inclusion and processing of new structures

To ease structure processing, a heuristics-based method was included in ULYSSES. This module checks for atomic clashes caused by overly-short bonds. Afterwards, the program goes over the atoms in the molecule and checks for the chemical neighbourhood. It then identifies certain patterns, which are associated with chemical groups and their properties. This is used to estimate the total molecular charge. We currently include several classes of functional groups, and more will be added in the future.

The program available from MISATO further includes basic electronic processing of the structures. This includes counting of electrons (with warnings issued if radicals are present), Frontier Molecular Orbital Analysis, bond-order calculation and AM1 charges. The latter may be directly input into programs like Amber.

**Tab. S3:** RMS of Koopman ionization potentials compared against experimental data, collected from CCCBDB [1]. RMS in eV. The ring type within quotation marks means that we only look at the ring extension. Thus for instance an “hexyl” includes benzene and cyclohexane.

| | n compound | AM1 | PM6 | B3LYP 6-311G** | M062X 6-311G** | MP2 6-311G** | CCSD(T) 6-311G** |
|---|---|---|---|---|---|---|---|
| Alkanes | 16 | 1.9 | 1.55 | 2.78 | 2.23 | 3.46 | 0.9 |
| Alkenes | 44 | 2.85 | 1.84 | 2.85 | 3.03 | 2.26 | 0.6 |
| Alkyne | 9 | 5.44 | 2.47 | 3.42 | 5.94 | 4.12 | 0.51 |
| Diene | 21 | 0.45 | 0.53 | 2.47 | 1.13 | 0.26 | --- |
| Alcohol | 16 | 6.23 | 4.69 | 4.75 | 6.61 | 5.9 | 1.36 |
| Ether | 17 | 0.57 | 1.89 | 2.86 | 1.11 | 1.39 | --- |
| Acid | 8 | 0.76 | 0.63 | 2.91 | 1.17 | 1.46 | --- |
| Ester | 8 | 0.85 | 0.48 | 2.95 | 0.87 | 1.42 | --- |
| Ketone | 8 | 3.49 | 3.04 | 3.41 | 1.05 | 3.88 | --- |
| Aldehyde | 11 | 8.48 | 6.68 | 6.03 | 7.49 | 8.47 | 1.01 |
| Nitroso | 10 | 7.75 | 5.52 | 6.68 | 8.21 | 9.46 | 15.5 |
| Nitro | 4 | 6.47 | 4.61 | 5.37 | 8.2 | 8 | 2.59 |
| Amide | 7 | 0.19 | 2.53 | 3 | 1.4 | 1.15 | --- |
| Halogen | 118 | 5.57 | 3.81 | 4.27 | 4.63 | 5.58 | 1.55 |
| Amine | 22 | 5.52 | 4.28 | 4.07 | 6.62 | 5.33 | 0.73 |
| $RCH_3$ | 128 | 3.98 | 2.68 | 3.43 | 4.33 | 3.77 | 1.06 |

| | | | | | | | |
|---|---|---|---|---|---|---|---|
| $RRCH_2$ | 110 | 1.43 | 0.99 | 2.85 | 2.02 | 1.73 | 1.07 |
| RRRCH | 22 | 0.69 | 0.46 | 2.73 | 1.03 | 1.11 | --- |
| RRRRC | 8 | 1.36 | 0.71 | 2.64 | 1 | 1.01 | --- |
| “Propyl” | 21 | 4.36 | 3.2 | 3.9 | 3.48 | 4.64 | 0.65 |
| “Butyl” | 9 | 0.65 | 0.59 | 2.6 | 1.02 | 0.81 | --- |
| “Pentyl” | 34 | 0.6 | 1.44 | 2.58 | 1.14 | 0.74 | --- |
| “Hexyl” | 27 | 2.75 | 0.53 | 2.48 | 4.27 | 0.75 | --- |
| Aromatic | 15 | 4.69 | 0.63 | 2.35 | 5.93 | 0.55 | --- |
| Bicycle | 5 | 0.81 | 0.92 | 2.48 | 1.07 | 0.34 | --- |
| Phenyls | 10 | 4.69 | 0.66 | 2.2 | 5.93 | 0.41 | --- |
